# The Virtual Brain links transcranial magnetic stimulation evoked potentials and inhibitory neurotransmitter changes in major depressive disorder

**DOI:** 10.1101/2024.11.25.622620

**Authors:** Timo Hofsähs, Marius Pille, Lucas Kern, Anuja Negi, Jil Mona Meier, Petra Ritter

**Author notes:** Equal contribution, co-last authors (Jil Mona Meier, Petra Ritter).

## Abstract

**Background:** Transcranial magnetic stimulation evoked potentials (TEPs) show promise as a biomarker in major depressive disorder (MDD), but the origin of the increased TEP amplitude in these patients remains unclear. Gamma aminobutyric acid (GABA) may be involved, as TEP peak amplitude is known to increase with GABAergic activity in healthy controls. We employed a computational modeling approach to investigate this phenomenon.

**Methods:** Whole-brain simulations in ‘The Virtual Brain’ (thevirtualbrain.org), employing the Jansen and Rit neural mass model, were optimized to simulate TEPs of healthy individuals (*N*_*subs*_=20, 14 females, 24.5±4.9 years). To mimic MDD-like impaired inhibition, a GABAergic deficit was introduced to the simulations by altering one of two selected inhibitory parameters, the inhibitory synaptic decay rate *b* or the number of inhibitory synapses *C*_4_. The TEP amplitude was quantified and compared for all simulations.

**Results:** The inhibitory synaptic decay rate showed a quadratic correlation (r=0.99, p<0.001) and the number of inhibitory synapses a negative exponential correlation (r=0.99, p<0.001) with the TEP amplitude. Moreover, significant correlations between these simulation-derived values and all TEP peaks and troughs were detected (p<0.001). Thus, under local parameter changes, we were able to alter the TEP amplitude towards pathological levels, i.e. creating an MDD-like increase of the global mean field amplitude in line with empirical results.

**Conclusions:** Our model suggests specific GABAergic deficits as the cause of increased TEP amplitude in MDD patients, which may serve as therapeutic targets. This work highlights the potential of whole-brain simulations in the investigation of neuropsychiatric diseases.

## Introduction

Investigative and therapeutic applications of transcranial magnetic stimulation (TMS) in major depressive disorder (MDD) show great, yet insufficiently utilized, potential (Farzan, 2024; Fitzsimmons et al., 2024; Gonsalves et al., 2024; van Rooij et al., 2024). Treatment success of repetitive TMS (rTMS) remains variable (Brunoni et al., 2017) and biomarkers lack understanding and standardization (Julkunen et al., 2022), fostered by the complexity and elusiveness of MDD pathomechanisms (Filatova et al., 2021). However, the recent success of accelerated rTMS treatment (Cole et al., 2022; Cole et al., 2020) and the potential diagnostic applications of TMS (Cao et al., 2021) in this high-burden disease (James et al., 2018; Vos et al., 2016; World Health Organization, 2022) encourage to deepen the knowledge of TMS effects in MDD patients, paving the way for its broader clinical use.

One piece of the puzzle is the unclear role of the main inhibitory neurotransmitter gamma aminobutyric acid (GABA) in TMS-evoked potentials (TEPs) of MDD patients (Dhami et al., 2020; Voineskos et al., 2019). The biomarker TEP measures cortical excitability and provides information about the excitatory-inhibitory balance, as its peaks and troughs emerge from summations of excitatory and inhibitory postsynaptic potentials (PSPs, (Hill et al., 2016; Tremblay et al., 2019)). Inhibition is disrupted in MDD patients, as the GABA hypothesis of depression claims (Luscher et al., 2011; Sanacora et al., 1999), which has been formulated as lower GABA levels were detected in MDD patients (Gerner & Hare, 1981; Godfrey et al., 2018; Hu et al., 2023; Petty & Sherman, 1984; Rajkowska et al., 2007). In healthy controls, higher GABA concentration measured by magnetic resonance spectroscopy (MRS, (Du et al., 2018)) and pharmacologically induced GABAergic inhibition (Darmani et al., 2016; Premoli et al., 2014) correlate with a higher amplitude of single-pulse TEP peaks. In MDD patients, the amplitude of TEP peaks is larger than in healthy controls (Dhami et al., 2020; Voineskos et al., 2019) and reduces with rTMS treatment (Biermann et al., 2022; Dhami et al., 2022; Strafella et al., 2023; Voineskos et al., 2021). From the current literature, the question how lower GABA levels can cause higher TEP amplitudes in MDD patients remains unanswered.

Whole-brain simulations are a versatile investigation tool to gain new insights into brain physiology and pathology, such as neurotransmitter mechanisms (Coronel-Oliveros, Cofré, et al., 2021; Deco, Cruzat, et al., 2018; Kringelbach et al., 2020; Naskar et al., 2021). The open-source software ‘The Virtual Brain’ (TVB, (Ritter et al., 2013; Sanz-Leon et al., 2015)) provides a framework to perform whole-brain simulations for various applications, e.g. to explore physiological brain-network mechanisms (Koller et al., 2024; Schirner et al., 2023), investigate disease pathologies (Courtiol et al., 2020; Falcon et al., 2015; Stefanovski et al., 2019), or simulate neuromodulation techniques (An et al., 2022; Kunze et al., 2016; Meier et al., 2022). Further, TVB is currently applied for the first time in a prospective clinical trial for presurgical planning in epilepsy patients (Wang et al., 2023). For our study, we make use of the advantages of TVB simulations, by manipulating GABA in individualized whole-brain models to generate an impaired inhibition as it occurs in MDD and dissect its effects on TEPs.

## Methods and Materials

### Data

An open-access dataset of concurrent electroencephalography (EEG) and TMS (Biabani C Rogasch, 2019) was employed to optimize the SCs to generate realistic TEPs of healthy subjects in the whole-brain simulations of this study. The dataset contains 62-channel EEG data of *N*_*subs*_=20 healthy subjects (24.5±4.9 years, 14 females) during single-pulse TMS of the left-hemispheric hand area of the primary motor cortex (M1), averaged over at least 100 repetitions with an intensity of 120% of the resting motor threshold. The empirical TEP timeseries of one example subject is shown in Figure 1F. The original study (Biabani et al., 2019) had been approved by the ethics committee of Monash University, Melbourne, Australia (Nr. CF15/822 - 2015000371). The further data processing for this study received approval by the ethics committee of Charité - Universitätsmedizin Berlin, Germany (Nr. EA4/045/24). A data sharing and processing agreement was set up and signed by the data controllers and processors of this study who worked with the data. For information about the preprocessing of the EEG data, please refer to the original publication (Biabani et al., 2019).

**Figure 1:**
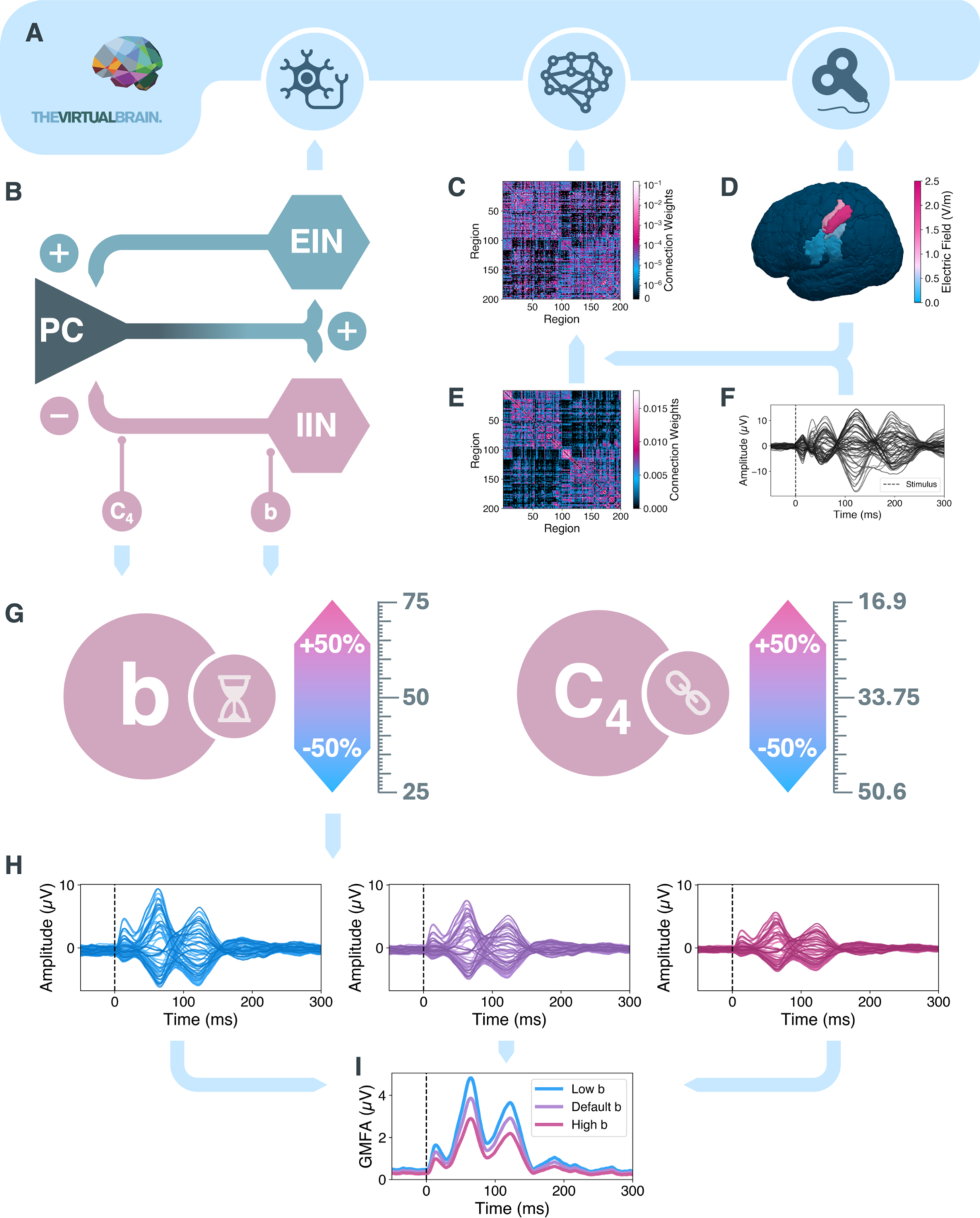
Workflow of the methodological steps. **A)** Whole-brain simulations with ‘The Virtual Brain’ were set up. **B)** Neural activity was generated by the Jansen and Rit neural mass model (Jansen & Rit, 1995). **C)** The connection weights between all regions were defined in the subject-specific optimized structural connectivity (SC) matrix *M* (depicted for example subject sub-15, diagonal set to 0 for visualization purposes). **D)** Amplitude of transcranial magnetic stimulation of left-hemispheric motor cortex per brain region on a surface brain. Five left-hemispheric regions received stimulation. **E)** Averaged and transformed empirical SC matrix *M͂*. To generate simulated transcranial magnetic stimulation evoked potential (TEP) with a high correlation to the empirical TEP **(F)**, the empirical SC matrix *M͂* was optimized, generating subject specific SC matrices *M* (C). **G)** Whole-brain simulations were repeated with either the inhibitory synaptic decay rate *b* or number of inhibitory synapses *C*_4_ altered to values in the range of −50% to +50% from their default values. **H)** Examples of simulated TEPs for low (left, blue), default (middle, purple) and high (right, magenta) inhibitory synaptic decay rate. **I)** The TEP amplitude was quantified by the GMFA. Stimulus onset is marked with a vertical black dashed line in panels (F), (H) and (I). EIN, excitatory interneuron population; GMFA, global mean field amplitude; IIN, inhibitory interneuron population; ms, milliseconds; µV, microvolt; PC, pyramidal cell population; V/m, volt per meter.

### Whole-Brain Simulations with The Virtual Brain

We employed TVB (version 2.7.1) to generate *in-silico* replications of the healthy subject empirical TEPs via whole-brain simulations ((Sanz Leon et al., 2013; Schirner et al., 2022), Figure 1A, Supplementary Section ‘Whole-Brain Simulations’, thevirtualbrain.org). Regional brain activity was simulated by one neural mass model per region (NMM, Figure 1B, Section ‘Neural Mass Model’). Interaction between all brain regions was defined by their connectome, consisting of region distances and structural connectivity (SC, Figure 1C, Supplementary Section ‘Connectome’). The whole-brain simulations were perturbed by virtual TMS (Figure 1D, Supplementary Section ‘Virtual TMS’). Simulated EEG timeseries were generated from the simulated source-level timeseries via a forward solution (Supplementary Section ‘Forward Solution’). SC matrices were optimized to generate simulated TEPs with a high correlation to empirical TEPs (Figure 1F) by employing a gradient-descent algorithm ((Momi et al., 2023), Figure 1C-F, Supplementary Section ‘Parameter Optimization’).

### Neural Mass Model

The activity of each brain region was simulated with the NMM by Jansen and Rit (JR, (Jansen & Rit, 1995)). Its ability to generate realistic electrophysiological data and TEPs made it an eligible choice for this work (Kunze et al., 2016; Momi et al., 2023; Spiegler et al., 2010). The JR model consists of three neuronal populations, including pyramidal cells (PC), excitatory interneurons (EIN) and inhibitory interneurons (IIN). The PC population is connected to the other populations by two separate feedback loops, while EIN and IIN do not interact directly (Figure 1B, Supplementary Figure S1). The activity of each population is described by the response function and the sigmoidal function. The response function, also known as the rate-to-potential operator, calculates the impulse response in

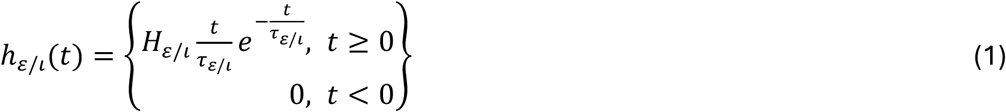

for the excitatory populations *ε*, i.e. PC and IIN, or the inhibitory population *l* (Supplementary Figure S2A), i.e. IIN. The function is defined by the maximum postsynaptic potential (PSP) *H*_*ε*/*l*_ and the synaptic time constant *τ*_*ε*/*l*_, the reciprocal of which represents ‘the lumped representation of […] passive membrane and all other spatially distributed delays in the dendritic network’ (Jansen & Rit, 1995). The response function can be expressed in two first-order linear inhomogeneous differential equations

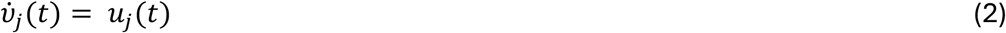

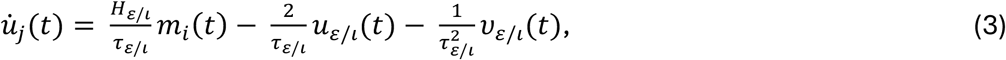

where the firing rate *m*_*i*_(*t*) of the presynaptic region *i* is applied as input to calculate the mean PSP *ν*_j_(*t*) of the postsynaptic population *j*. The sigmoid function S (Supplementary Figure S2B)

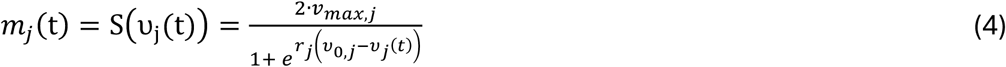

is known as the potential-to-rate operator and calculates the firing rate *m*_*j*_(*t*) of a neuronal population *j* at time point *t* depending on its mean PSP *ν*_*j*_ = ∑_*i*_ *ν*_*ij*_. Parameter *ν*_*max,j*_ is half of the maximum of the mean firing rate of population *j* in spikes per second, *ν*_0,*j*_ shows the firing threshold in millivolt for which a 50% firing rate is achieved in population *j* and the steepness *r*_*j*_ of the curve at the inflection point defines the variability of firing threshold values within the NMM for population *j* (Spiegler et al., 2010). The framework in Equations 1-4 is adapted to each of the three neural populations, resulting in six differential equations. For the NMM in region *k*, the PSP *y*_0,*k*_(*t*) of PC is calculated by

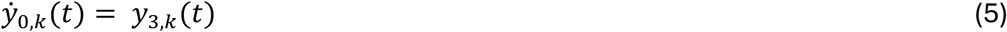

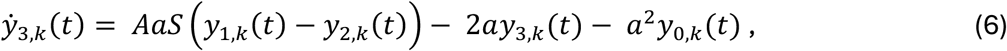

the PSP *y*_1,*k*_(*t*) of EIN with

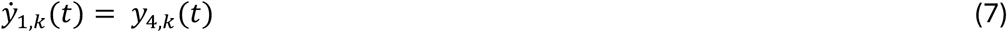

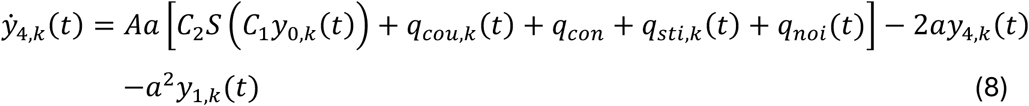

the PSP *y*_2,*k*_(*t*) of IIN in

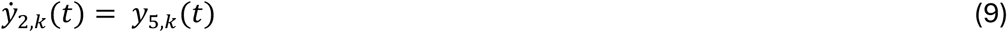

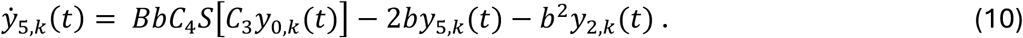

The general term *H*_*ε*/*l*_ is defined as *A* in the excitatory populations *ε*, i.e. PC and EIN, and *B* in the inhibitory population *ι*, i.e. IIN. Equivalently, the reciprocals of *τ*_*ε*/*l*_ are defined as *a* and *b*, the excitatory and inhibitory synaptic decay rates, which refers to the rate at which PSPs rise and decay. Parameters *C*_1–4_ refer to the number of synapses between the neural populations, *C*_1_ for PC to EIN, *C*_2_ for EIN to PC, *C*_3_ for PC to IIN and *C*_4_ for IIN to PC. In Equation 8, which generates the PSP of the EIN, several terms *q* for additional input are added, including coupled activity from other regions *q*_*cou*,*k*_(*t*), constant external input *q*_*con*_, external perturbation *q*_*sti*,*k*_(*t*), i.e. TMS, and noise *q*_*noi*_(*t*). The transformed coupled activity from other regions *q*_*cou*,*k*_(*t*) for region *k* is defined in another sigmoid function

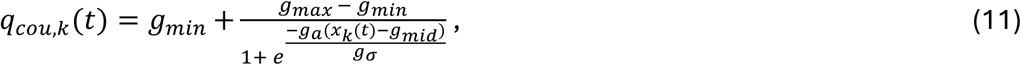

in which *g*_*min*_ and *g*_*max*_ define the lower and upper saturation values, *g*_*mid*_ defines the midpoint, *g*_σ_ the steepness, and *g*_*a*_ the scaling factor. The factor *x*_*k*_(*t*) is the summed long-range coupled activity input before transformation through the sigmoid in Equation 11 from all other regions *l* into region *k* at time point *t* and is defined as

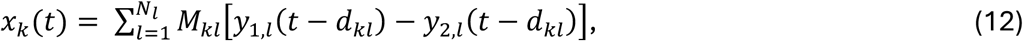

where the PSP per region *l* is calculated by the difference of state variables *y*_1,*l*_ and *y*_2,*l*_ at time point *t* minus the delay *d*_*kl*_ between regions *k* and *l*, derived by their distance divided by the conduction speed (2.5m/s). The activity is scaled by the individual connection strength between regions *k* and *l*, as defined in the optimized SC matrix *M* (Figure 1C, Supplementary Section ‘Parameter Optimization’) and summed up over all regions *N*_*l*_=200 as input for region *k*. The values for all described parameters are listed in Supplementary Table S1.

### Parameter Alteration

In a first step, and before parameter alterations, parameter optimization of SC weights was performed *N*_*reps*_=100 times per subject, resulting in *N*_*opt*_=2,000 optimized parameter sets (Supplementary Section ‘Optimizing Structural Connectivity’). Secondly, after optimizing the SC and simulating healthy control TEPs in TVB utilizing the optimized SC, specific parameters within the whole-brain simulations were altered to introduce GABAergic deficits and assess the impact on TEPs. TEP simulations in TVB were repeated with changed values of *b* or *C*_4_ (Supplementary Section ‘Inhibitory Feedback Loop Analysis’). One of the two parameters was altered to values of −50% to +50% of the respective default values in steps of 2% (*b*: default=50, range=[25,75]; C_4_: default=33.75, range=[16.875,50.625], Figure 1G-H), creating a total of *N*_*alt*_=101 simulations (50 altered values per parameter plus 1 default simulation) per optimized parameter set. Whenever one parameter was altered, the other one was kept at default value. The effect of the maximum and minimum alterations of *b* and C_4_ on the JR-inherent functions are shown in Supplementary Figure S2 (A: *b* alteration effects on the response function; B: C_4_ alteration effects on sigmoidal function). For each of the simulations, the global mean field amplitude (GMFA) was calculated to quantify the TEP amplitude (Figure 1I, Supplementary Section ‘TEP Amplitude Quantification’). The relative GMFA was calculated by normalizing the GMFA values from all simulations with altered *b* and C_4_ to the GMFA from the simulation with default *b* and C_4_ for the same individual, and expressing the result as a percentage. For each *b* and *C*_4_ value the mean over all *N*_*reps*_=100 per subject was calculated to generate the relative GMFA per subject. A supplementary analysis per subject has been performed (Supplementary Section ‘Impact of *b* and *C*_4_ alterations per subject’). However, for the major analysis, the mean over the *N*_*subs*_=20 values was taken, resulting in *N*_*alt*_=101 independent mean relative GMFA values, 50 for *b*, 50 for *C*_4_ and one default value. Following regressions were performed over the 50 altered values of *b* plus one default, and one over the 50 altered values of *C*_4_ plus one default, separately.

### Statistics

To assess the relationship between the mean GMFA and *b* / *C*_4_ value, we compared three types of regression models, including a linear, quadratic and an exponential model. For each model, the fit was quantified by the sum of squared errors (SSE) between the regression model and the mean GMFA, and the model with the lowest SSE was selected for representation. For the linear model, regression parameters were estimated using ordinary least squares, and significance was evaluated by a Wald test. The quadratic polynomial model was likewise fitted by ordinary least squares, with overall model significance assessed by an F-test. For the exponential model, parameters were estimated by nonlinear least-squares optimization, with standard errors derived from the covariance matrix and p-values computed from the corresponding t-statistics. A significance threshold of p<0.05 was applied for all tests.

## Results

An average correlation over all *N*_*subs*_=20 subjects and *N*_*reps*_=100 optimization repetitions between the simulated and empirical TEP timeseries of a Pearson correlation coefficient (PCC) =0.633 was achieved.

The relation of JR parameter *b* with the relative GMFA of the TEP is depicted in Figure 2. The inhibitory synaptic decay rate *b* shows a significant quadratic relation to the relative GMFA of the overall TEP (Figure 2A, r=0.99, p<0.001). The average relative GMFA increases for values above and below the default inhibitory synaptic decay rate. A quadratic correlation has been detected in all individual subjects (Supplementary Figure S4). While relative GMFA increases with decreasing inhibitory synaptic decay rate values in all subjects, the behavior for values above default differs from strong GMFA increase to slight GMFA decrease.

**Figure 2:**
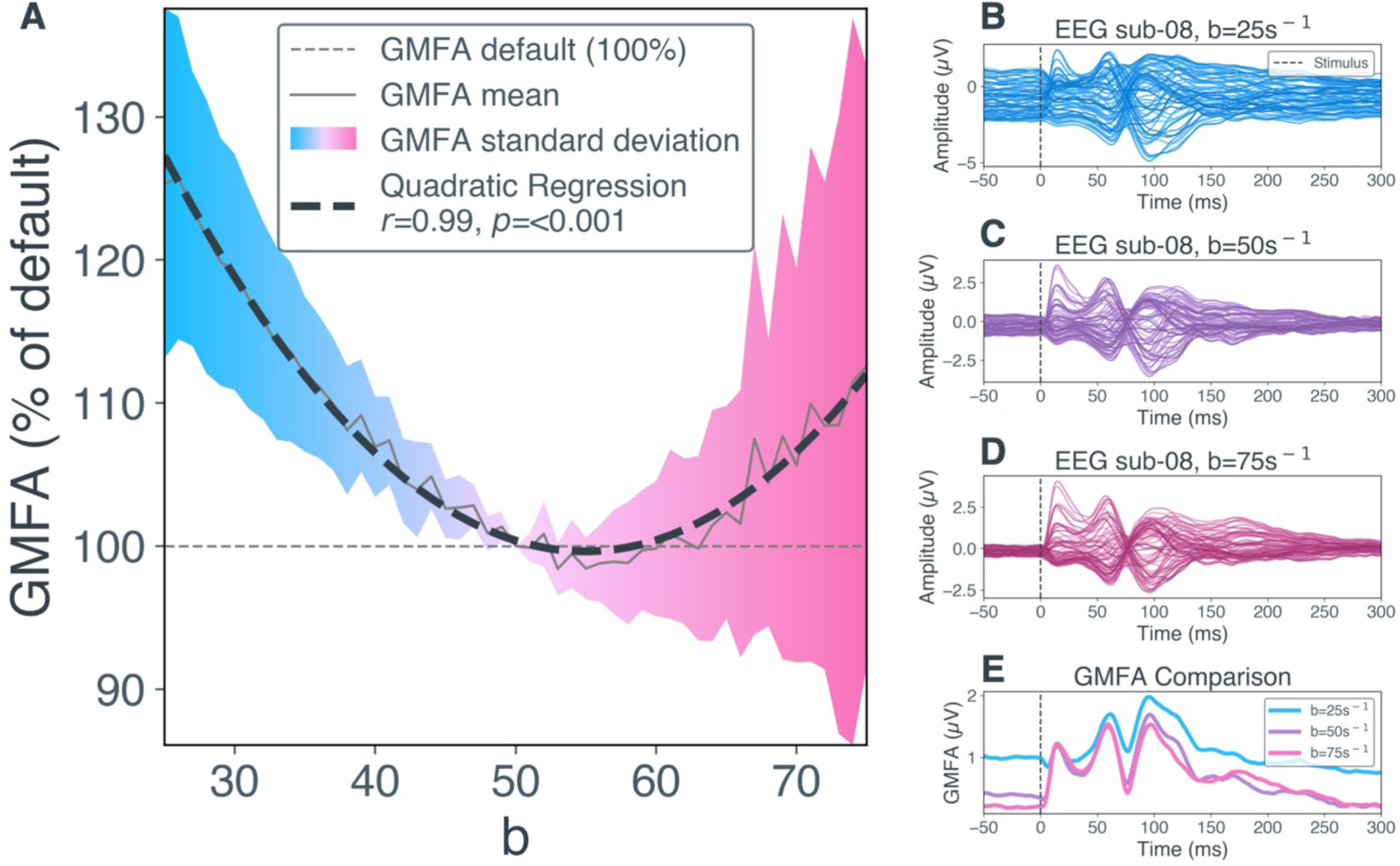
Effects of inhibitory synaptic decay rate *b* on global mean field amplitude (GMFA) of transcranial magnetic stimulation evoked potentials (TEPs). **A)** The relative GMFA amplitude (in % relative to GMFA at default parameter value *b*=50) per parameter value for *b* and quadratic regression is shown here. The solid grey line shows the mean relative GMFA over *N*_*opt*_=2,000 optimized parameter sets. The dashed black line shows the quadratic regression calculated over these mean values. The colors indicate the standard deviation of relative GMFA values, i.e. blue for −50% to default and magenta for default to +50%. The dashed grey line shows GMFA at default parameter value *b*=50. **B)** EEG timeseries of example subject sub-08 from a simulation with low inhibitory synaptic decay rate (*b*=25). Each blue line shows the activity measured in one EEG channel. The dashed black line indicates stimulus onset at 0ms. **C)** EEG timeseries of sub-08 from a simulation with default inhibitory synaptic decay rate (*b*=50). **D)** EEG timeseries of sub-08 from a simulation with high inhibitory synaptic decay rate (*b*=75). **E)** Comparison of GMFA (i.e. area under the curve, summed up over all channels in **B-D**) from low (*b*=25, blue), default (*b*=50, purple) and high (*b*=75, magenta) inhibitory synaptic decay rate simulations for sub-08. EEG, electroencephalography; GMFA, global mean field amplitude; ms, milliseconds; µV, microvolt; s, second; sub, subject.

For the number of inhibitory synapses C_4_, a significant exponential relation to the relative GMFA of the overall TEP has been detected (Figure 3A, r=0.99, p<0.001). For values below the default number of inhibitory synapses, the average relative GMFA increases exponentially, whereas for values above the default it remained relatively stable, asymptotically approaching a level slightly below 100% of the default. Exponential regression showed the best fit in ten subjects, while quadratic correlation showed the best fit in the remaining ten subjects (Supplementary Figure S5). For the number of synapses from PC to IIN C_3_, a significant quadratic correlation to the relative GMFA of the overall TEP has been shown (Supplementary Section ‘Alterations of *C*_3_ ‘).

**Figure 3:**
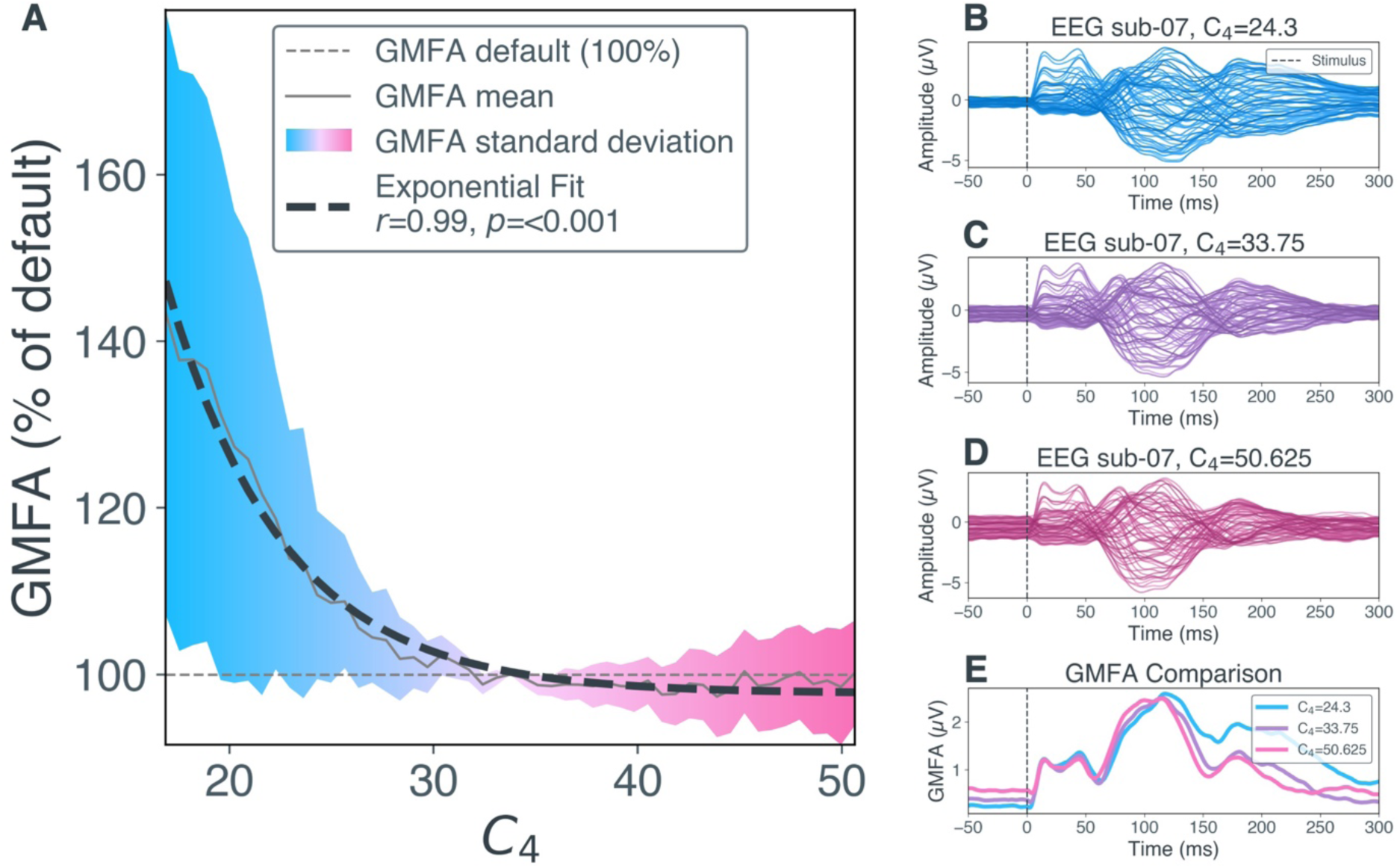
Effects of number of inhibitory synapses *C*_4_ on global mean field amplitude (GMFA) of transcranial magnetic stimulation evoked potentials (TEPs). **A)** The relative GMFA amplitude (in % relative to GMFA at default parameter value *C*_4_=33.75) per parameter value for *C*_4_ and exponential regression is shown here. The solid grey line shows the mean relative GMFA over *N*_*opt*_=2,000 optimized parameter sets. The dashed black line shows the exponential regression calculated over these mean values. The colors indicate the standard deviation of relative GMFA values, i.e. blue for −50% to default and magenta for default to +50%. The dashed grey line shows GMFA at default parameter value *C*_4_=33.75. **B)** EEG timeseries of example subject sub-07 from a simulation with low number of inhibitory synapses (*C*_4_=24.3). Each blue line shows the activity measured in one EEG channel. The dashed black line indicates stimulus onset at 0ms. **C)** EEG timeseries of sub-07 from a simulation with default number of inhibitory synapses (*C*_4_=33.75). **D)** EEG timeseries of sub-07 from a simulation with high number of inhibitory synapses (*C*_4_=50.625). **E)** Comparison of GMFA (i.e. area under the curve, summed up over all channels in **B-D**) from low (*C*_4_=24.3, blue), default (*C*_4_=33.75, purple) and high (*C*_4_=50.625, magenta) inhibitory synaptic decay rate simulations for sub-08. EEG, electroencephalography; GMFA, global mean field amplitude; ms, milliseconds; µV, microvolt; s, second; sub, subject.

Figure 4 shows a significant quadratic relation between GABA function and TEP amplitude for all analyzed peaks and throughs, with the single exception of *C*_4_ in P185 (Figure 4B4) that demonstrated a higher goodness-of-fit level with an exponential function.

**Figure 4:**
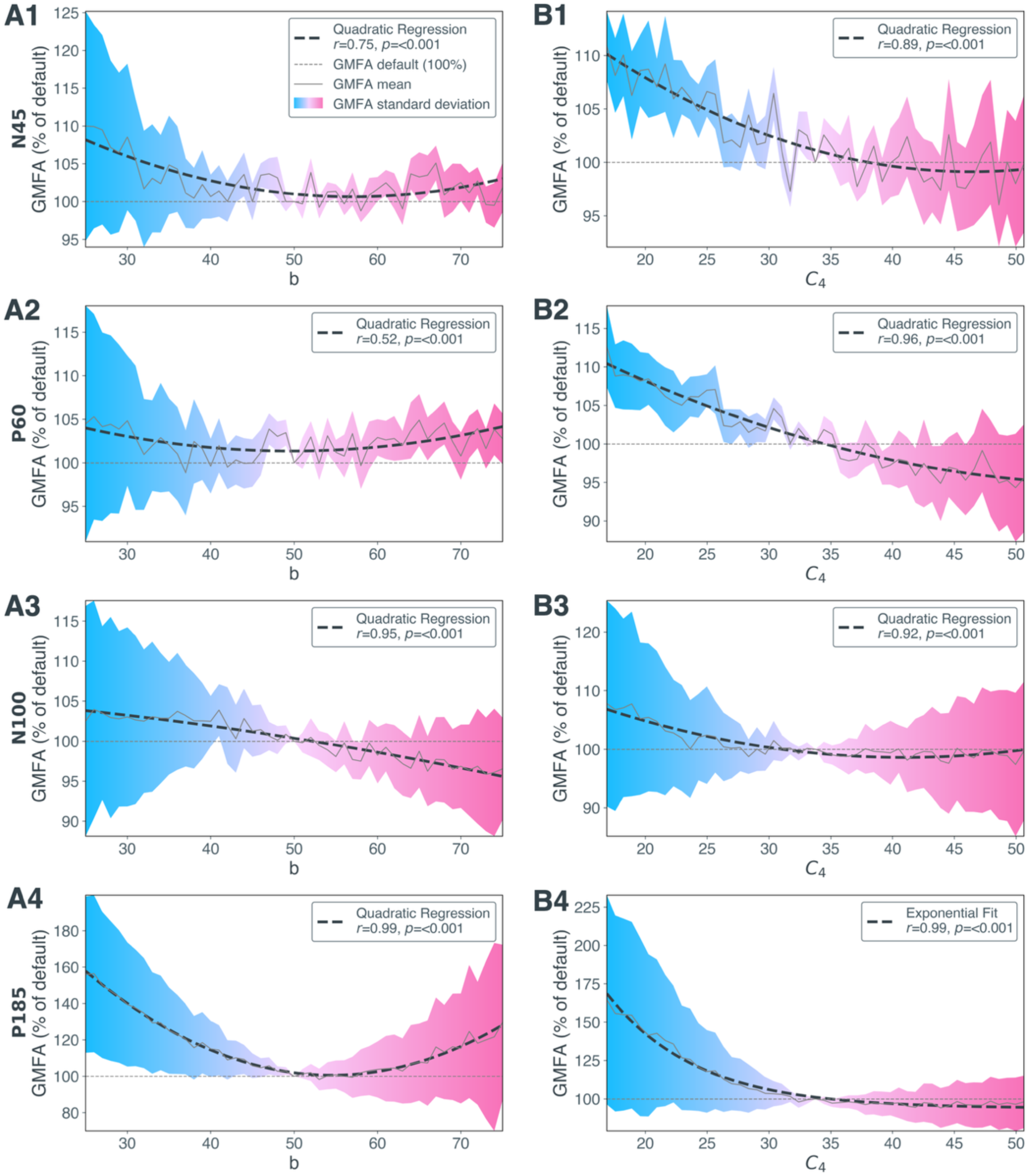
Relative global mean field amplitude (GMFA) changes due to alterations of inhibitory synaptic decay rate *b* and number of inhibitory synapses *C*_4_ per transcranial magnetic stimulation (TMS) evoked potential (TEP) peak N45, P60, N100 and P185. The relative GMFA amplitude (in % relative to GMFA at default parameter values *b*=50 **(A1-A4)** and *C*_4_=33.75 **(B1-B4)**, default is depicted as grey dotted line) per parameter values for *b* **(A1-A4)** and *C*_4_ **(B1-B4)** and quadratic or exponential regression are shown here. The solid grey line shows the mean relative GMFA over *N*_*opt*_=2,000 optimized parameter sets, the dashed black line shows the optimal regression calculated over these mean values. The colors indicate the standard deviation of relative GMFA values, i.e. blue for −50% to default and magenta for default to +50%. GMFA, global mean field amplitude.

## Discussion

In this study, we investigated the role of GABAergic neurotransmission in the genesis of TEP alterations with a whole-brain simulation approach. Our model shows a high correlation between the simulated and empirical TEPs. We were able to alter the TEP amplitude in silico by adjusting the inhibitory synaptic decay rate and the number of inhibitory synapses.

We reproduced an empirical phenomenon with computational modeling. TEP changes were created through simulation of GABAergic deficits. Significant GMFA alterations were achieved in every subject individually as well as in the overall average. In a previous study, the average GMFA of TEPs in MDD patients was approximately 130% of the one in healthy controls (Voineskos et al., 2019). In our model, comparable overall TEP amplitude alterations were achieved via lowering *b* from 50 to 26 (125.73% of the GMFA at default settings) and *C*_4_ from 33.75 to 19.575 (131.25%). Thus, these parameter values represent the most realistic mimicking of MDD pathology in our model. Furthermore, GMFA increase was shown empirically for all single TEP peaks and troughs in MDD patients (Dhami et al., 2020; Voineskos et al., 2019). In line with this, we detected a significant correlation for all peak’s and trough’s GMFA with both parameter alterations. Similar to the overall TEP, the largest GMFA increase was detected for the lowest parameter values, confirming the previously described trend. In line with empirical findings in (Dhami et al., 2020), the late TEP components show the largest alterations. Our findings were achieved in a theory-driven approach by single parameter manipulations, following previous modeling approaches (Cabrera-Álvarez et al., 2024; Stefanovski et al., 2019). This work contributes to the growing number of studies investigating the mechanisms of MDD and possible therapeutic targets through computational whole-brain modeling (An et al., 2022; Deco et al., 2024; Escrichs et al., 2024; Wang et al., 2022).

The results of this study propose a mechanistic connection between two phenomena occurring in MDD patients, GABAergic deficits (Luscher et al., 2011; Sanacora et al., 1999) and an increased TEP amplitude (Dhami et al., 2020; Voineskos et al., 2019). We drew this mechanistic connection by observing TEP amplitude increases after mimicking GABAergic deficits through alterations of local inhibitory model parameters. Thereby, we establish a possible explanation for the previously described open question regarding the mismatch of GABA levels (meaning the concentration of the molecule measured in blood plasma, cerebrospinal fluid or MRS) and GABAergic inhibition (meaning the power of inhibition measured, for example via paired-pulse TMS) in MDD patients. A comparable disconnection of GABA levels and GABAergic inhibition has previously been reported in healthy controls, as GABA levels assessed with MRS did not correlate with paired-pulse TMS measures in these subjects (Cuypers & Marsman, 2021). The authors suggested that the methods might measure distinct parts of the GABAergic neurotransmission (Cuypers & Marsman, 2021). MRS, for example, mainly measures intracellular concentrations of GABA (Godfrey et al., 2018; Gonsalves et al., 2022), while paired-pulse TMS measures capture GABA_A_- and GABA_B_-receptor dependent function (Cuypers & Marsman, 2021). In addition to the existing range of research methods, our *in-silico* approach allows the targeted modulation of GABAergic inhibition, independent of pharmacological or paired-pulse TMS approaches. In our simulations, we implemented two separate mechanisms to model reduced GABAergic inhibition, by reducing the inhibitory synaptic decay rate or the number of inhibitory synapses, both leading to increased TEP amplitudes. Hence, our findings appear in contrast with the theory that increased TEP amplitudes in MDD patients arise from an increased postsynaptic GABA turnover (i.e. higher GABAergic inhibition but lower GABA levels), which has been postulated in the empirical trials showing increased TEP amplitudes in MDD (Dhami et al., 2020; Voineskos et al., 2019). The TEP amplitude increase might rather arise from higher cortical excitability through lower GABAergic inhibition, which is disconnected from the relationship of GABA and single TEP peaks.

According to our results, the first mechanism that might cause TEP changes is the disruption of the excitation-inhibition (E/I) timing through alteration of the inhibitory synaptic decay rate. More specifically, we demonstrated a quadratic relationship between this parameter value and the TEP amplitude, i.e. by decreasing or increasing the inhibitory synaptic decay rate, the average TEP amplitude increased. The inhibitory synaptic decay rate reflects the aggregate temporal dynamics of inhibitory signaling at the population level. The parameter encompasses all biological processes that affect the time course of an action potential within the dendritic network of the IIN, from its arousal at the dendrite, until GABA reached its receptors in the post-synapse and later its re-uptake into the GABAergic neuron (Jansen & Rit, 1995). Previous studies showed that the processes most crucial for the overall time are the calcium ion dynamics, which evoke the IIN action potential to cause GABA release (Sabatini & Regehr, 1996; Schneggenburger & Neher, 2000), and the SNARE complex, which fuses the transport vesicle with the IIN cell membrane to release GABA into the postsynaptic cleft (Acuna et al., 2014). GABAergic inhibition physiologically modulates brain activity by determining time windows that allow excitation by inhibiting the postsynaptic membrane potential (Pelkey et al., 2017). The TEP captures the physiological, well-timed balance between excitation and inhibition, because it arises from distinct spatiotemporal cascades of excitatory and inhibitory PSPs (Hill et al., 2016; Rogasch & Fitzgerald, 2013; Tremblay et al., 2019). The previously described empirical factors contributing to the inhibitory neurotransmission speed may therefore be involved in MDD pathology and explain both the GABAergic deficit and the increased TEP amplitude.

Our results reveal a second possible explanation of TEP amplitude increases in MDD, which is a decreased number of inhibitory synapses. By lowering the number of inhibitory synapses, a shifted E/I balance with an overweight of excitation is created, causing a TEP amplitude increase. Similar results were achieved via decreasing the number of excitatory synapses which activate the inhibitory feedback loop (Supplementary Figure S6). In line with these findings, a previous study showed an increase in the average PC membrane potential after impairment of the number of inhibitory synapses in a single node model (Wendling et al., 2002). The mathematical parameter for the number of inhibitory synapses may biologically be interpreted as all factors contributing to the hyperpolarizing strength of the IIN on the PC, including the total number of synapses of the IIN onto the PC, the number of functioning postsynaptic GABA_A_- and GABA_B_-receptors, the level of GABA concentration in the synaptic cleft, and the postsynaptic ion influx (Pelkey et al., 2017; Schousboe & Waagepetersen, 2017). Decreased GABA_A_- and GABA_B_-receptor mediated activity has been found in MDD patients in multiple clinical trials (Bajbouj et al., 2006; Duman et al., 2019; Levinson et al., 2010; Sun et al., 2016). More generally, an increase in cortical excitability has been shown in MDD patients (Kinjo et al., 2021), and previous results have indicated that the vulnerability of GABAergic neurotransmission by IIN might be the main factor disrupting the E/I balance in MDD (Duman et al., 2019). Our results support these findings and highlight molecular interactions involved in the hyperpolarization strength of IIN within the complex mechanism of inhibition in MDD.

This study contributes to the growing body of literature that is suggesting the GABAergic deficit as a potential target mechanism for MDD therapy (Möhler, 2012). Significantly lower GABA concentrations in MDD patients compared to healthy control were found in currently depressed MDD patients, but not in currently remitted MDD patients, suggesting a link between the GABA concentration and depression symptoms (Godfrey et al., 2018). Increases in GABA levels were observed following the application of selective serotonin reuptake inhibitors in healthy (Bhagwagar et al., 2004) and depressed subjects (Sanacora et al., 2002), and after electroconvulsive therapy in MDD patients (Sanacora et al., 2003). Moreover, symptom improvement after rTMS therapy was correlated with an increase in MRS-detected GABA concentration (Gonsalves et al., 2022) and rTMS therapy has been associated with a decrease in the amplitude of single TEP peaks in multiple studies (Biermann et al., 2022; Dhami et al., 2022; Strafella et al., 2023; Voineskos et al., 2021). Hence, literature emphasizes the involvement of GABA in the MDD pathology and its restoration by treatment. In line with this, we created digital brain twins of healthy subjects and mimicked a GABAergic deficit through two previously described mechanisms. The model suggests that GABAergic neurotransmission can be targeted to ‘reverse’ a depressive state to a healthy state, i.e. in terms of a normalization of the TEP amplitude. Corresponding to this, GABA has previously been identified as suitable target for MDD therapy (Möhler, 2012). Novel therapy approaches impacting GABAergic neurotransmission are benzodiazepines, positive allosteric GABA-receptor modulators, ketamine, and rTMS (Duman et al., 2019; Fogaça & Duman, 2019; Gerhard et al., 2020; Krystal et al., 2023). Benzodiazepines act via allosteric binding of the postsynaptic GABA_A_ receptor, which strengthens the GABAergic postsynaptic hyperpolarization (Griffin et al., 2013). Strengthening the GABAergic postsynaptic hyperpolarization was modeled in our simulations by increasing the number of inhibitory synapses. Therefore, the model cannot only be used to draw conclusions about the mechanisms of MDD but provides suggestions for treatment as well. In fact, none of the treatment approaches impacting the GABAergic neurotransmission are currently part of the first-line therapy guidelines in MDD (American Psychological Association, 2019), but rTMS has been brought up as a possible first-line approach in MDD, given its promising results and the low success rates of current first-line therapies (O’Sullivan et al., 2024). The results of this study emphasize the potential of tackling the GABAergic deficit for an effective treatment of MDD.

From a dynamical systems perspective, we modeled an emergent network phenomenon by showing the sensitivity of simulated TEPs to local inhibitory parameter alterations. The impact of local inhibitory parameters of the JR model onto neural dynamics in single node non-perturbed models has previously been shown (Spiegler et al., 2010; Wendling et al., 2002). On a network scale, the alterations of local JR parameters can be utilized to investigate complex phenomena, such as integration-segregation (Coronel-Oliveros, Castro, et al., 2021; Coronel-Oliveros et al., 2023; Coronel-Oliveros et al., 2024) or dementia (Cabrera-Álvarez et al., 2024; Coronel-Oliveros et al., 2024; Stefanovski et al., 2019; Triebkorn et al., 2022). In addition, perturbation analysis, in which a system is perturbed by an external stimulus to observe the following effect, is an established method for examining global network dynamics (Deco, Cabral, et al., 2018; Gollo et al., 2017). In the simplified two-node network, perturbation-induced dynamics in the JR model are primarily determined by the connectivity strength between the nodes (Ahmadizadeh et al., 2018; Basu et al., 2018; Grimbert & Faugeras, 2006; Jansen & Rit, 1995). Similarly, the response evoked by TMS *in-vivo* is shaped by the network properties, as the activity propagates through diverse pathways over the whole cortex (Farzan et al., 2016). Recent work has modeled these complex network responses to TMS perturbation in healthy subjects (Momi et al., 2023). In our work, we combined the perturbational approach with local inhibitory JR parameter alterations. The correlation between empirical and simulated TEPs was achieved through SC optimization in our model, confirming the crucial role of the network in the response to TMS perturbation. Moreover, single local inhibitory JR parameter alterations created global alterations in our whole-brain simulations, mimicking a phenomenon empirically observed in MDD patients. Interestingly, alterations of the two local parameters simultaneously showed nonlinear behavior, as the strongest GMFA increase was detected via increasing one parameter and decreasing the other (Supplementary Section ‘Simultaneous Alterations of *b* and *C*_4_’). Further work is necessary to exploit the great potential of nonlinear dynamics in whole-brain simulations for the diagnosis and treatment of MDD using TMS.

While this study provides valuable insights, we acknowledge the following limitations. First, the proposed model was not validated with empirical MDD patient data, since, to the best of our knowledge, no suitable open-access dataset was available and attempts to receive non-open data were unsuccessful. A validation against patient data would increase our model’s reliability, ideally by linking simulation parameters to depression symptoms. However, our mimicked MDD TEPs were quantitatively compared with empirical published summary results of MDD patients. The link between our simulated impairment of inhibition, increased TEP amplitudes and patient symptoms would address another limitation: Without validation with patient data, the lowering of *b* and *C*_4_ may be interpreted differently than as MDD pathology, since GABAergic deficits can also be caused by other reasons. Additionally, our model focuses specifically on the GABA hypothesis of depression, excluding other relevant hypotheses, such as the monoamine or inflammatory hypotheses (Filatova et al., 2021; Pitsillou et al., 2020), which future work utilizing other specified NMM could investigate. Furthermore, whole-brain models are a simplification of reality, omitting specific biological aspects. For example, local parameters were equal in all regions and simulations, despite neurotransmitter variations across the cortex in reality, and the GABAergic changes were introduced homogeneously over the whole cortex. Furthermore, the Schaefer parcellation does not contain subcortical regions. Spatial details of TMS on a sub-regional level were not considered due to averaging the electric field per region. External input into the neural mass model, including TMS stimulus, global coupling input and noise, was added to the excitatory population activity only (Equation 8) to drive pyramidal cell activity. Moreover, TMS was applied to M1 and not the dorsolateral prefrontal cortex, the area commonly targeted in MDD treatment (Lefaucheur et al., 2020). We chose M1 as stimulation target, because we decided to develop an already validated model further, which was created with the utilized dataset (Biabani & Rogasch, 2019) and optimization method (Momi et al., 2023), and induced GABAergic deficits to model MDD. Interestingly, recent empirical results had proposed M1 as a suitable target for rTMS treatment for MDD, suggesting mechanisms comparable to the dorsolateral prefrontal cortex (Hu et al., 2024). Lastly, while TEP is a promising biomarker, it has faced criticism due to potential artifacts (Conde et al., 2019) and the lack of standardized procedures (Belardinelli et al., 2019).

Neither MDD (Pitsillou et al., 2020), nor TMS-EEG (Farzan, 2024), nor GABA in psychiatric disorders (Fee et al., 2017; Fries et al., 2023), are fully understood today. Generally, whole-brain simulations are suitable for targeting open research questions of this intricately linked nature and high complexity level. Additionally, these simulations provide a suitable framework for personalization, a goal shared by psychiatric research in general (Wium-Andersen et al., 2017), and TMS research in particular (Cocchi & Zalesky, 2018; Klooster et al., 2021; Modak & Fitzgerald, 2021). Regarding our specific model, future efforts should address the previously described limitations and first and foremost incorporate MDD patient data. Establishing a link between MDD symptoms and GABAergic alterations could provide additional validity to the model. Another improvement would be the inclusion of a plasticity component (Jannati et al., 2023) to simulate disease progression and investigate the mechanisms of successful therapy as previously proposed (Wilson et al., 2018; Wilson et al., 2016). Further personalization could be achieved by integrating individual connectomes (Meier et al., 2024; Triebkorn et al., 2024), spatial sub-regional TMS distribution (Thielscher et al., 2015), and heterogeneous neurotransmitter maps (Deco, Cruzat, et al., 2018) via heterogeneous variation of local parameters. Interestingly, individual subjects exhibit behavior that runs counter to the average. Further research could clarify whether these individual models could be utilized to elaborate the subject-specific variability of TEPs and the differences in therapeutic success of TMS. Ultimately, the goal is to enable pre-therapeutic simulations of MDD patients’ brains to optimize stimulation parameters for rTMS treatment.

In this study, we explored the role of GABAergic neurotransmission in the mechanisms underlying increased TEP amplitudes, comparable to those occurring in MDD patients. We optimized whole-brain simulations to generate TEPs of healthy subjects and then modified inhibitory simulation parameters to replicate the GABAergic deficits in MDD. Our findings revealed that lowering the inhibitory synaptic decay rate *b* and the number of inhibitory synapses C_4_ led to an increased TEP amplitude. Therefore, this work suggests a mechanistic explanation for TEP alterations in MDD through two separate avenues of altered local inhibition. Future development of novel treatment approaches could consider these pathomechanisms as potential targets. By showcasing our personalized whole-brain modeling approach in this study, our results present a steppingstone towards personalized modeling-enriched MDD therapy in the future.

## Acknowledgements

PR acknowledges support by the Virtual Research Environment at the Charité Berlin – a node of EBRAINS Health Data Cloud, by EU Horizon Europe program Horizon EBRAINS2.0 (101147319), Virtual Brain Twin (101137289), EBRAINS-PREP 101079717, AISN – 101057655, EBRAIN-Health 101058516, Digital Europe TEF-Health 101100700, EU H2020 Virtual Brain Cloud 826421, Human Brain Project SGA2 785907; Human Brain Project SGA3 945539, German Research Foundation SFB 1436 (project ID 425899996); SFB 1315 (project ID 327654276); SFB 936 (project ID 178316478; SFB-TRR 295 (project ID 424778381); SPP Computational Connectomics RI 2073/6-1, RI 2073/10-2, RI 2073/9-1; DFG Clinical Research Group BECAUSE-Y 504745852, PHRASE Horizon EIC grant 101058240; Berlin Institute of Health and Foundation Charité. JM acknowledges funding by the Deutsche Forschungsgemeinschaft (DFG, German Research Foundation) – Project-ID 424778381 – TRR 295.

This manuscript was previously uploaded as a preprint on *biorxiv* on November 25, 2024 (DOI: 10.1101/2024.11.25.622620).

TH and JM acknowledge the use of the large language model ‘ChatGPT’ (chatgpt.com/) for corrections and improvement of semantics of already written text during the writing process of this script.

## Contributions

- Conceptualization: TH, JM, PR;
- Data Curation: TH, JM;
- Formal Analysis: TH;
- Funding Acquisition: PR
- Investigation: TH, LK;
- Methodology: TH, MP, JM, AN;
- Project Administration: JM, PR;
- Resources: PR;
- Software: TH, MP, PR;
- Supervision: JM, PR;
- Validation: TH, JM;
- Visualization: TH;
- Writing – Original Draft: TH;
- Writing – Review C Editing: TH, JM, MP, PR, LK, AN;

We thank current and former members of the ‘Brain Simulation Section’ at Charité Universitätsmedizin, Berlin, Germany, Halgurd Taher, Jan Stasiński and Chanida Chayopathum for insightful discussions over the course of this project. Computation has been performed on the High-Performance Cluster for Research and Clinic of the Berlin Institute of Health, Berlin, Germany.

## Data and Code Availability

The empirical TEP dataset (Biabani & Rogasch, 2019) is publicly available for download (bridges.monash.edu/articles/dataset/TEPs-_SEPs/7440713). The entire code to generate the results presented in this study is openly available (github.com/virtual-twin/TMS_MDD/).

