## Supplemental Information for "The Virtual Brain links transcranial magnetic stimulation evoked potentials and inhibitory neurotransmitter changes in major depressive disorder"

### Whole-Brain Simulations

The initial conditions for the neural mass model (NMM) state variables (Equation 5-10) were randomly chosen from a uniform distribution between 0 and 1. To solve the differential equations, a stochastic extension of the Euler integrator (Sanz-Leon et al., 2015) was chosen with a step size of 0.25ms. White noise with a variance of $\sigma_{noi}$=0.0003 was added to the fifth state variable $y_{4,k}$ of the NMM for all regions $k$ (Equation 8). The parameter optimization (Supplementary Section 'Parameter Optimization') was repeated $N_{rep}$=100 times per subject to increase the robustness of the results, creating $N_{opt}$=2,000 optimized parameter sets. For each of these sets, simulations with parameters altered were performed ($N_{alt}$=101, Method Section 'Parameter Alteration'), leading to a total of $N_{sim}$=$N_{opt} \cdot N_{alt}$ =202,000 simulations. All computations were performed on a high-performance cluster.

#### *Connectome*

The connectome utilized in our whole-brain simulations was defined by two matrices, the distance matrix and the structural connectivity (SC) matrix. The distance matrix contains the distance in millimeters between all regions in the connectome and was defined by the white-matter fiber tract lengths between regions. The SC defines the strength of connection between all regions in the connectome and was based on the number of streamlines between regions. Both matrices utilized in this work, the distance matrix and the SC matrix, were derived from diffusion-weighted imaging and structural magnetic resonance imaging data from the 'Human Connectome Project' (Van Essen et al., 2013). Both matrices were averaged over 400 healthy subjects (21-35 years, 230 females) and parcellated into $N_l$=200 regions according to the Schaefer atlas (Schaefer et al., 2017). The connectome data was taken from Momi et al. ((Momi et al., 2023), github.com/GriffithsLab/PyTepFit/tree/main). For details regarding the preprocessing of the empirical data underlying the connectome, please refer to the original publication (Momi et al., 2023). Before usage in our whole-brain simulations, the SC underwent a transformation and a subject-specific parameter optimization. Transforming the original averaged streamline-based empirical SC matrix $\bar{M}$ using the natural logarithm and Frobenius norm in

$$\widetilde{M}_{kl} = \frac{\ln(1+\bar{M}_{kl})}{\sqrt{\sum_{\delta=1}^{N_l}\sum_{\varepsilon=1}^{N_l}[\ln(1+\bar{M}_{\delta\varepsilon})]^2}} \tag{S1}$$

for the connections between all regions $k$ and $l$ for $N_l$=200 created an averaged transformed SC matrix $\widetilde{M}$ (Figure 1E). Following thereafter, $\widetilde{M}$ was numerically altered by taking the negative of its Laplacian, a method previously described in literature (Abdelnour et al., 2018; Atasoy et al., 2016; Raj et al., 2022). The outcome was optimized subject specifically to generate matrix $M$ (Figure 1C, Supplementary Section 'Parameter Optimization'), which was then utilized in the whole-brain simulations.

#### *Virtual TMS*

The same spatiotemporal stimulus settings as in Momi et al. (Momi et al., 2023) were applied to simulate transcranial magnetic stimulation (TMS, Figure 1D). The open-source software 'SimNIBS' (Thielscher et al., 2015) was employed to approximate the spatial distribution of TMS on left-hemispheric motor cortex (M1). The software generated a surface-based estimation of an electric field (e-field), which was downsampled to the Schaefer atlas (Schaefer et al., 2017) by assigning each vertex to a region and calculating the average e-field strength over all vertices per region. However, not the whole e-field but only an area underneath the TMS coil center surpasses the empirically-derived threshold to evoke action potentials in the underlying neural tissue (Romero et al., 2019). This threshold is approximated in empirical settings by determining the resting motor threshold (rMT), i.e. the minimum stimulation amplitude at which at least 50% of the pulses elicit a motor-evoked potential of ≥50μV in the hand muscle the stimulated cortical area is innervating (Klomjai et al., 2015). The TMS amplitude was set to 120% rMT in the empirical trial (Biabani et al., 2019). To transform the e-field according to Romero et al. (Romero et al., 2019), the highest value in the simulated e-field array was therefore regarded as equal to 120% stimulation amplitude, and all regions receiving an estimated e-field strength below rMT (i.e. 83% (=100%/120%) of the maximum value in the e-field array) were set to zero to simulate TMS. The e-field strength was below the rMT of 83% of the maximum in 195 of 200 regions, leaving five regions located around the left-hemispheric M1 receiving virtual TMS input in our simulations (Figure 1D). The stimulus is applied in the simulation by adding the input variable $q_{sti,k}(t)$ to the excitatory population state variable in the respective region $k$, as in the original NMM publication ((Jansen & Rit, 1995), Equation 8, Supplementary Figure S1). The stimulation was applied for 10ms after 1000ms to avoid the initial simulation transient. The total duration per simulation was 1400ms. In Figure 1H, Figure 2B-E and Figure 3B-E only the time frame 950-1300ms is depicted and the stimulus onset at 1000ms is denoted as 0ms for visualization purposes.

#### *Forward Solution*

Sensor-level EEG timeseries for whole-brain simulations $E_{sim}$ were calculated as

$$E_{sim} = P \cdot L \,, \qquad \text{(S2)}$$

i.e. by multiplication of the simulated source-level activity $P$ with a leadfield matrix $L$. The matrix $P$ contains the source level activity per region for the simulation duration. The source level activity defined as difference between state variables $y_{1,k} - y_{2,k}$ for each region $k$ following David et al. (David et al., 2006). The leadfield matrix $L$ was taken from Momi et al. ((Momi et al., 2023), github.com/GriffithsLab/PyTepFit/tree/main).

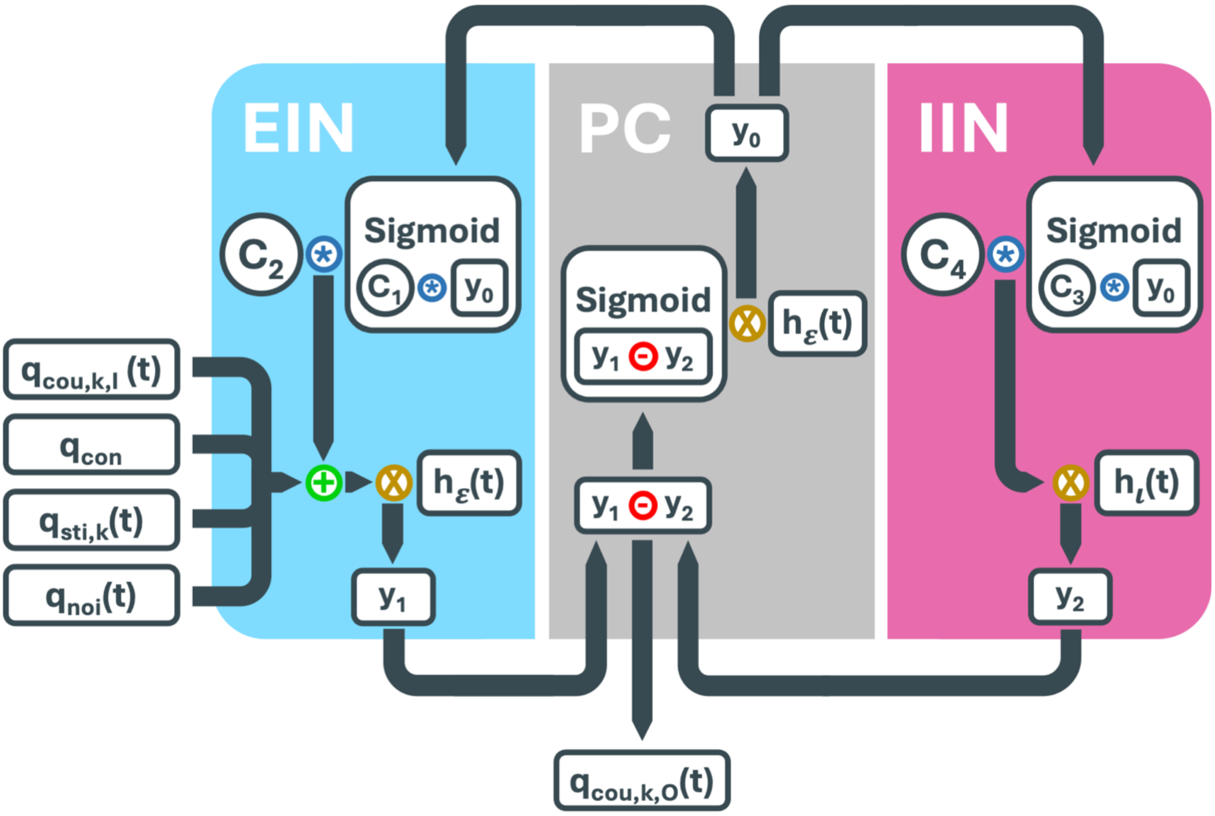


**Supplementary Figure S1: Schematic overview of the Jansen and Rit neural mass model ((Jansen & Rit, 1995), Equations 1-12).** The three neuronal populations, pyramidal cells (PC, state variable $y_0$), excitatory interneurons (EIN, state variable $y_1$) and inhibitory interneurons (IIN, state variable $y_2$), are connected in two separate feedback loops. The activity of each population is generated by a set of equations, including the sigmoid function and the response function $h$ for excitatory $\varepsilon$ and inhibitory $\iota$ populations. The input and the output of the sigmoidal function is multiplied by synapse scaling factors ($C_1/C_2$ in EIN; $C_3/C_4$ in IIN). External input into the region $k$ is added to the EIN and consists of input global coupling $q_{cou,k,I}(t)$ from other regions into region $k$, constant input $q_{con}$, external input by stimulation (i.e. TMS) $q_{sti,k}(t)$ and noise $q_{noi}(t)$. The activity generated in region $k$ affects other regions via output global coupling $q_{cou,k,O}(t)$.

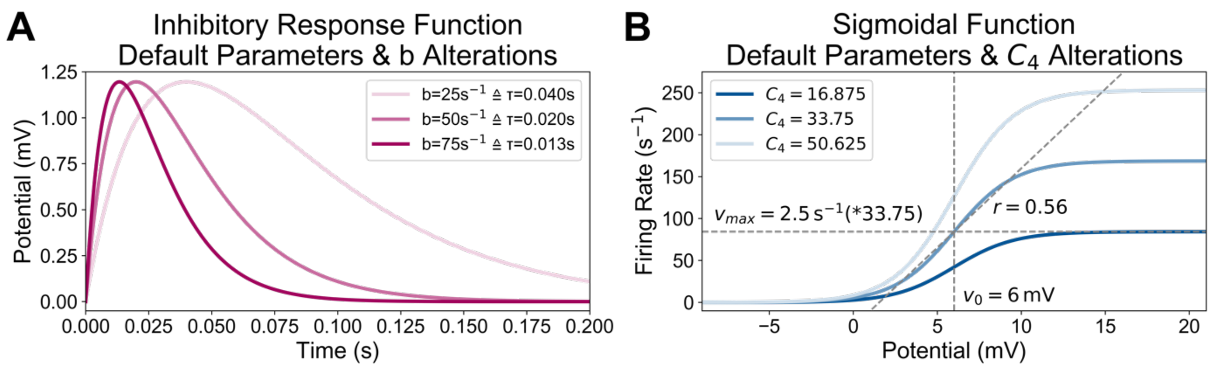


**Supplementary Figure S2: Graphical depiction of the two core functions of the Jansen and Rit neural mass model ((Jansen & Rit, 1995), Equations 1-12) - inhibitory response function (A) and sigmoidal function (B) - and the effect of parameter alterations on them. A):** The response function is also known as rate-to-potential operator (Equations 1-4). The inhibitory response function calculated with default parameters, including the inhibitory synaptic decay rate $b$=50s$^{-1}$ is depicted in middle magenta. The effects of alterations of $b$ to -50% ($b$ =25s$^{-1}$, light magenta) and +50% ($b$ =75s$^{-1}$, dark magenta) of its default value are shown as well. The higher $b$ is, the faster the membrane potential reaches its maximum. **B)** The sigmoidal function is also known as potential-to-rate operator (Equation 4). Input is the membrane potential of the respective neural population and output is the corresponding firing rate in spikes per second. The function is defined by the firing threshold $v_0$=6mV, half of the maximum mean firing rate $v_{max}$=2.5s$^{-1}$ of the population, and the steepness of the curve $r$=0.56. The sigmoidal function in the inhibitory feedback loop is scaled by the number of inhibitory synapses $C_4$=33.75 (default) and shown in middle blue. The effects of its alterations to -50% ($C_4$=16.875, light blue) and +50% ($C_4$=50.625, dark blue) of its default value are depicted. The higher $C_4$ is, the higher is the possible maximum firing rate.

**Supplementary Table S1: Whole-brain simulation and optimization parameters.**

| Name | Value | Unit | Description |
|---|---|---|---|
| *Structural Connectivity* | | | |
| $\bar{M}$ | - | - | original streamline-based empirical SC matrix |
| $\widetilde{M}$ | - | - | transformed SC matrix |
| $M$ | - | - | optimized, subject-specific SC matrix |
| *Sigmoidal Coupling Parameters* | | | |
| $g_{min}$ | -0.5 | - | minimum of the sigmoidal coupling function |
| $g_{max}$ | 0.5 | - | maximum of the sigmoidal coupling function |
| $g_{mid}$ | 0 | - | midpoint of the sigmoidal coupling function |
| $g_a$ | 4 | - | scaling factor of sigmoidal coupling function |
| $g_\sigma$ | 1 | - | standard deviation of the sigmoidal coupling function |
| *Jansen & Rit Neural Mass Model Parameters* | | | |
| $A$ | 22 | mV | maximum amplitude of the EPSP (PC, EIN) |
| $B$ | 3.25 | mV | maximum amplitude of the IPSP (IIN) |
| $a$ | 100 | $s^{-1}$ | excitatory synaptic decay rate (PC, EIN)<br>(excitatory synaptic time constant inverse) |
| $b$ | 50 | $s^{-1}$ | default inhibitory synaptic decay rate (IIN)<br>(default inhibitory synaptic time constant inverse) |
| | 25 | $s^{-1}$ | lower bound of altered values for $b$ |
| | 75 | $s^{-1}$ | upper bound of altered values for $b$ |
| $C_1$ | 135 | - | number of synapses from PC to EIN |
| $C_2$ | 108 | - | number of synapses from EIN to PC |
| $C_3$ | 33.75 | - | number of synapses from PC to IIN |
| $C_4$ | 33.75 | - | number of synapses from IIN to PC (default value) |
| | 16.875 | - | lower bound of altered values for $C_4$ |
| | 50.625 | - | upper bound of altered values for $C_4$ |
| $v_{max}$ | 2.5 | $s^{-1}$ | half of the maximum firing rate of all neuronal populations |
| $v_0$ | 6 | mV | PSP for which half of the maximum firing rate of the neuronal population is achieved; can be interpreted as excitability of all neuronal populations |
| $r$ | 0.56 | $mV^{-1}$ | steepness at the firing threshold;<br>represents the variance of firing thresholds within the NMM |
| $q_{con}$ | 1e-9 | - | constant input |
| *Optimization Parameters* | | | |
| $\mu_p$ | $vec(\widetilde{M})$ | - | default value of parameters included in optimization,<br>$vec(\widetilde{M})$: vectorized upper triangle of the matrix $\widetilde{M}$ |
| $\sigma_M$ | 1000 | - | weighting of SC in optimization algorithm |
| *Quantities* | | | |
| $N_l$ | 200 | - | Number of brain regions in simulations (Schaefer parcellation, (Schaefer et al., 2017)) |
| $N_p$ | 13,861 | - | Number of connections per SC matrix in parameter optimization |

| | | | |
|---|---|---|---|
| $N_z$ | 62 | - | Number of channels on the EEG cap |
| $N_{subs}$ | 20 | - | Number of healthy subjects in empirical TEP dataset |
| $N_{reps}$ | 100 | - | Number of repetitions of the parameter optimization per subject |
| $N_{opt}$ | 2,000 | - | Number of optimized parameter sets ($N_{subs}$ x $N_{reps}$) |
| $N_{alt}$ | 101 | - | Number of parameter alterations per optimized parameter set |
| $N_{sim}$ | 202,000 | - | Total number of simulations performed for this study ($N_{opt}$ x $N_{alt}$) |

EFL, excitatory feedback loop; EIN, excitatory interneuron population; EPSP, excitatory postsynaptic potential; IFL, inhibitory feedback loop; IIN, inhibitory interneuron population; IPSP, inhibitory postsynaptic potential; m/s, meters per second; ms, milliseconds; mV, millivolt; NMM, neural mass model; PC, pyramidal cell population; PSP, postsynaptic potential; SC, structural connectivity; TEP, transcranial magnetic stimulation evoked potential.

### Parameter Optimization

#### *Algorithm*

The optimization of whole-brain simulation parameters to generate realistic TMS-evoked potentials (TEP) *in silico* was performed by adapting the method outlined by Momi et al. (Momi et al., 2023). The underlying optimizer is a gradient-descent algorithm that was recently proposed (Griffiths et al., 2022). All equations of the whole-brain simulation and parameter optimization are embedded in 'PyTorch' (Version 2.3.0), a machine-learning software package (Paszke et al., 2019) for building and training artificial neural networks. The PyTorch-inherent automatic differentiation capabilities and optimization algorithms allow for optimizing whole-brain simulation parameters to replicate electrophysiological data. The gradient-based 'ADAM' algorithm (Kingma & Ba, 2014) was deployed to find the optimal parameter constellation to minimize the cost function

$$\lambda = \beta + \gamma, \tag{S3}$$

where $\beta$ is the regularization penalty and $\gamma$ is the loss. The regularization penalty

$$\beta = \sum_{p=1}^{N_p} \frac{(\theta_p - \mu_p)^2}{\sigma_p} \tag{S4}$$

is summed up over all $N_p$=13,861 (Supplementary Section 'Optimizing Structural Connectivity') optimizing parameters $p$ and depends on their respective initial values $\mu_p$ (before optimization) and their individual parameter weightings $\sigma_p$. The term $\theta_p$ shows the altered value of parameter $p$. The mean-squared error loss function calculates the match between the currently simulated TEP in EEG timeseries $E_{sim,z}$ and empirical TEP in EEG timeseries $E_{emp,z}$ in

$$\gamma = \sum_{t=-100}^{300} \left[\frac{1}{N_z} \sum_{z=1}^{N_z} \left(E_{sim,z}(t) - E_{emp,z}(t)\right)^2\right] \tag{S5}$$

using a mean-squared-error loss function per time point $t$ in the time frame -100ms to 300ms when stimulus onset is regarded as 0ms and for the step size of 1ms and aggregated over all channels $z$. Following the method by Momi et al. (Momi et al., 2023), the similarity between empirical and simulated EEG was assessed with a second metric, the Pearson correlation coefficient (PCC), which is calculated in

$$\mathrm{PCC} = \frac{1}{N_z} \sum_{z=1}^{N_z} \left[ \frac{\sum_{t=-100}^{300} \left(E_{sim,z}(t) - \bar{E}_{sim,z}\right)\left(E_{emp,z}(t) - \bar{E}_{emp,z}\right)}{\sqrt{\sum_{t=-100}^{300} \left(E_{sim,z}(t) - \bar{E}_{sim,z}\right)^2 \sum_{t=-100}^{300} \left(E_{emp,z}(t) - \bar{E}_{emp,z}\right)^2}} \right] \quad \text{(S6)}$$

for the same time frame as in Equation S5. The parameter $\bar{E}$ is the average per channel $z$ over the same time frame of 100ms before to 300ms after stimulus onset.

#### *Optimizing Structural Connectivity*

Only the SC matrix was included in the parameter optimization, because a high correlation between empirical and simulated TEPs was achieved (average PCC=0.633). Hereby, every non-zero connection weight was optimized individually. Of the $N_l(N_l - 1)/2$=19,900 connections in the upper triangle of the original SC matrix, 6,039 were equal to zero and omitted by the optimization algorithm. Therefore, a total of $N_p$=13,861 connections were optimized for each individual (Figure 1C-F). The value of the transformed SC matrix $\widetilde{M}$ (Figure 1C, Supplementary Section 'Connectome') was chosen as default value $\mu_M$ (Equation S4) for all connections in all subjects. To restrict the parameters, the weighting $\sigma$ was set equally for all optimized parameters $p$ (i.e. all non-zero connections within $\widetilde{M}$) to $\sigma_p$=$\sigma_M$=1000 (Equation S4).

The optimization was performed $N_{reps}$=100 times per subject to mitigate the influence of random initial conditions, noise, and variability in optimization outcomes. Repeating $N_{reps}$=100 times on $N_{subs}$=20 subjects created $N_{opt}$=2,000 optimized parameter sets.

#### TEP Amplitude Quantification

To quantify the global TEP amplitude, the global mean field amplitude (GMFA, (Komssi & Kähkönen, 2006; Lehmann & Skrandies, 1980)) was calculated for all simulations. The GMFA $G$

$$G(t) = \sqrt{\frac{1}{N_z}\left[\sum_{z=1}^{N_z}(E_z(t) - E_{mean}(t))^2\right]} \tag{S7}$$

at time point $t$ is quantified with the EEG amplitude $E_z$ of channel $z$, the mean EEG amplitude $E_{mean}$ over all channels, and the total number of channels $N_z = 62$ (Figure 1I). The GMFA of the overall TEP was calculated for the time frame 55-275ms post-stimulus, for the N45 peak 45ms post-stimulus ±10ms, for P60 peak 60ms post-stimulus ±10ms, for N100 peak 100ms post-stimulus ±20ms and for P185 peak 185ms post-stimulus ±20ms, following earlier TEP studies (Dhami et al., 2020; Voineskos et al., 2019).

### Inhibitory Feedback Loop Analysis

Possible candidates that could serve as biological proxies of gamma aminobutyric acid (GABA) were identified among the mathematical whole-brain simulation parameters. Since GABA is the main inhibitory neurotransmitter of the human brain (Petroff, 2002), all JR inhibitory feedback loop parameters $B$, $b$, $C_3$ and $C_4$ (Equation 10, Supplementary Figure S1, Supplementary Table S1) were considered. However, since the parameters $B$ and $C_4$ have equivalent scaling effects, investigating both would be redundant (Equation 8). Compared to $B$, which represents the biologically plausible maximum value of the IPSP with a unit, i.e. 22mV (van Rotterdam et al., 1982), parameter $C_4$ is a proportional value, i.e. the average number of inhibitory synapses compared to the other synapse parameters $C_1$, $C_2$ and $C_3$. The more abstract character of $C_4$ compared to $B$ allows for stronger deviations from default without violating biological plausibility. Consequently, $B$ was excluded from the alterations. Furthermore, $C_3$ is the number of synapses between the pyramidal cell population and the inhibitory interneuron population and therefore represents glutamatergic excitatory neurotransmission. Due to the focus of this work on GABAergic neurotransmission, $C_3$ was excluded from the parameter alterations in the main analysis as well, leaving the inhibitory synaptic decay rate $b$ and the number of inhibitory synapses $C_4$ as the parameters to explore and model GABA in this study. The alteration of the parameter $C_3$ was performed in a separate analysis (Supplementary Information Section 'Alteration of $C_3$').

### Simultaneous Alterations of $b$ and $C_4$

In an additional analysis, the effect of simultaneous alterations of $b$ and $C_4$ on the TEP amplitude was investigated. Both parameters were altered between -50% and +50% of their default values in 12.5% steps, creating nine values per parameter and a grid of 81 variations in total. Simulations of all subjects and ten optimized parameter sets per subject were run with all 81 parameter constellations, generating 16,200 simulations in total (20 subjects x 10 optimization repetitions x 81 parameter variations). Mean relative GMFA per subject was computed similarly to the main results (Section 'Parameter Alteration'). The mean relative GMFA over all subjects was calculated and depicted in Supplementary Figure S3.

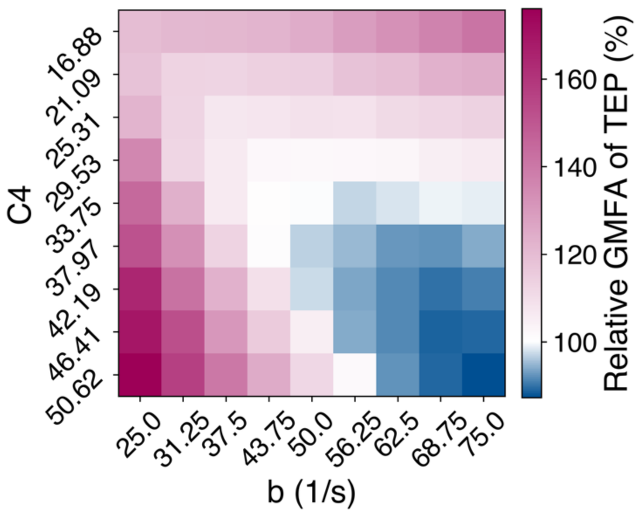


**Supplementary Figure S3: Effects of simultaneous alterations of inhibitory synaptic decay rate $b$ and number of inhibitory synapses $C_4$ on GMFA of TEPs.** Simulations of 20 subjects and ten optimized parameter sets were repeated with altered values of $b$ and $C_4$ in 81 parameter constellations. Mean relative GMFA was first calculated per subject over the ten optimized parameter sets. Secondly, mean relative GMFA over all subjects was calculated. Default parameter simulations are depicted in the central panel ($b$ =50/s, $C_4$=33.75). All other simulations show the relative GMFA, meaning the GMFA change in comparison to default. Blue shadings show TEP amplitude decrease, magenta shadings show TEP amplitude increase.

#### Impact of $b$ and $C_4$ alterations per subject

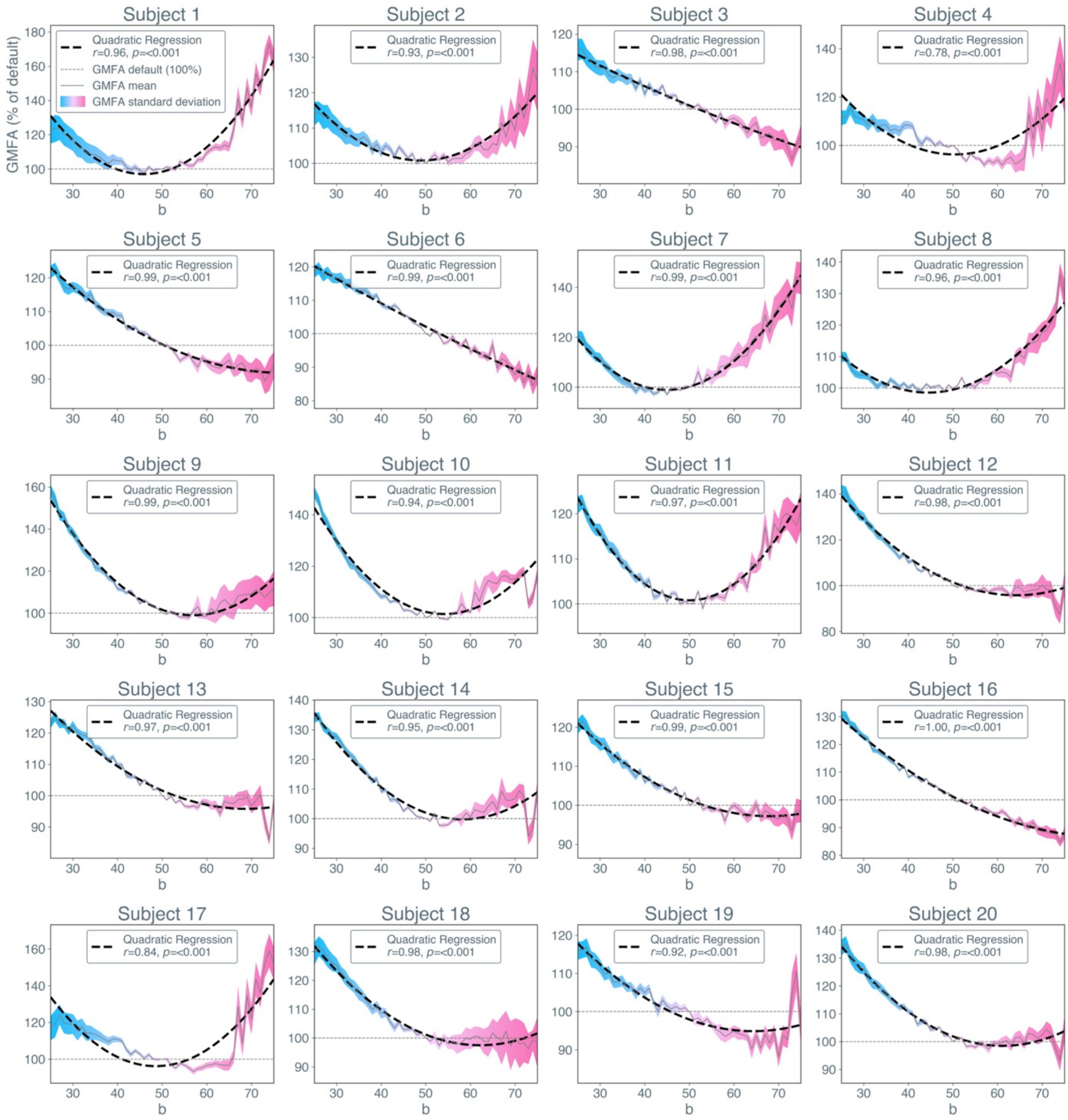


**Supplementary Figure S4: Effects of inhibitory synaptic decay rate $b$ alterations on relative global mean field amplitude (GMFA) of transcranial magnetic stimulation evoked potentials per subject.** The relative GMFA amplitude (in % relative to GMFA at default parameter value $b$=50, default is depicted as grey dotted line) per parameter value for $b$ and best fitting regression is shown here. The solid grey line shows the mean relative GMFA over $N_{reps}$=100 optimized parameter sets, the dashed black line shows the quadradic regression calculated over these mean values. The colors indicate the standard deviation of relative GMFA values, i.e. blue for -50% to default and magenta for default to +50%.

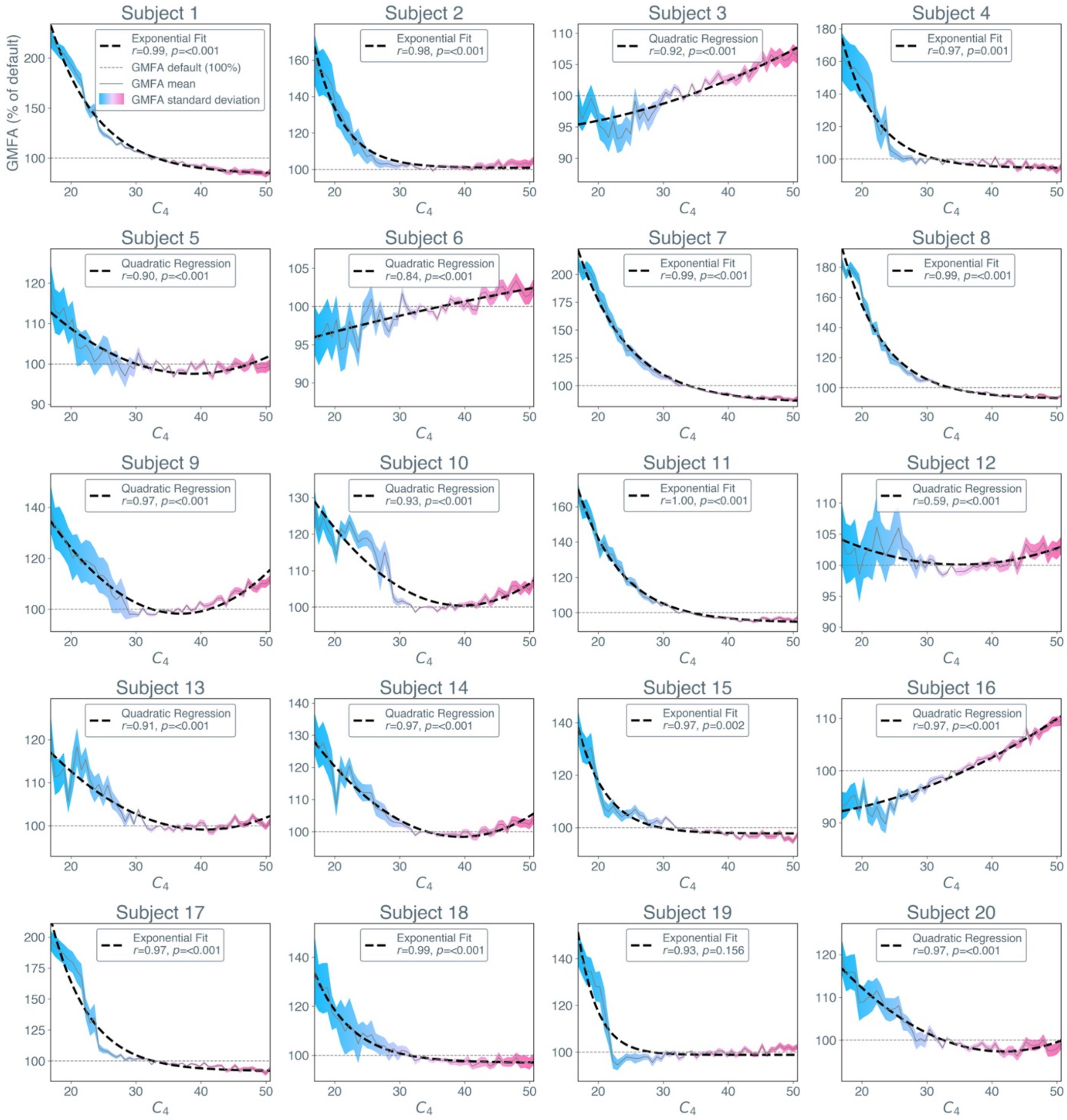


**Supplementary Figure S5: Effects of number of inhibitory synapses $C_4$ alterations on relative global mean field amplitude (GMFA) of transcranial magnetic stimulation evoked potentials per subject.** The relative GMFA amplitude (in % relative to GMFA at default parameter value $C_4$=33,75, default is depicted as grey dotted line) per parameter value for $C_4$ and best fitting regression is shown here. The solid grey line shows the mean relative GMFA over $N_{reps}$=100 optimized parameter sets, the dashed black line shows the quadradic regression calculated over these mean values. The colors indicate the standard deviation of relative GMFA values, i.e. blue for -50% to default and magenta for default to +50%.

### Alterations of $C_3$

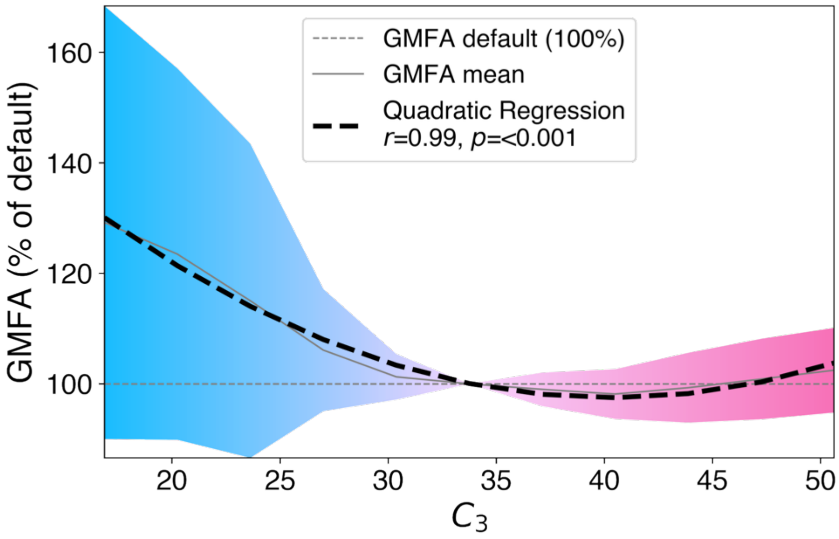


**Figure S6: Effects of number of synapses between pyramidal cells and excitatory interneurons $C_3$ on GMFA of transcranial magnetic stimulation evoked potentials.** The relative GMFA amplitude (in %, relative to GMFA at default parameter value $C_3$=33.75, default is depicted as light grey dashed line) per parameter value for $C_3$ and linear as well as quadratic regression is shown here. $C_3$ was altered from -50 to +50% in 10% steps. Alterations of $C_3$ were applied to simulations of all 20 subjects and ten optimized parameter sets per subject. The solid gray line shows the mean relative GMFA over 200 optimized parameter sets. The black dotted line shows the linear regression calculated over these mean values. The black dashed line shows the quadratic regression calculated over these mean values. The colors indicate the standard deviation of relative GMFA values, i.e. blue for -50% to default and magenta for default to +50%.
